# Force Field Dependent DNA Breathing Dynamics: A Case Study of Hoogsteen Base Pairing in A6-DNA

**DOI:** 10.1101/2022.05.04.490579

**Authors:** Sharon Emily Stone, Dhiman Ray, Ioan Andricioaei

## Abstract

The Hoogsteen (HG) base pairing (bp) conformation, commonly observed in damaged and mutated DNA helices, facilitates DNA repair and DNA recognition. The free energy difference between HG and Watson-Crick (WC) base pairs has been computed in previous studies. However, the mechanism of the conformational transition is not well understood. A detailed understanding of the process of WC to HG base pair transition can provide deeper understanding of DNA repair and recognition. In an earlier study, we explored the free energy landscape for this process using extensive computer simulation with the CHARMM36 force field. In this work, we study the impact of force field models in describing the WC to HG base pairing transition using meta-eABF enhanced sampling, quasi-harmonic entropy calculation, and non-bonded energy analysis. The secondary structures of both base pairing forms and the topology of the free energy landscapes were consistent over different force field models, although the relative free energy, entropy and the interaction energies tend to vary. The relative stability of the WC and HG conformations is dictated by a delicate balance between the enthalpic stabilization and the reduced entropy of the structurally rigid HG structure. These findings highlight the impact that subtleties in force field models can have on accurately modeling DNA base pair dynamics and should stimulate further computational investigations into other dynamically important motions in DNA.

**Graphical TOC Entry:** 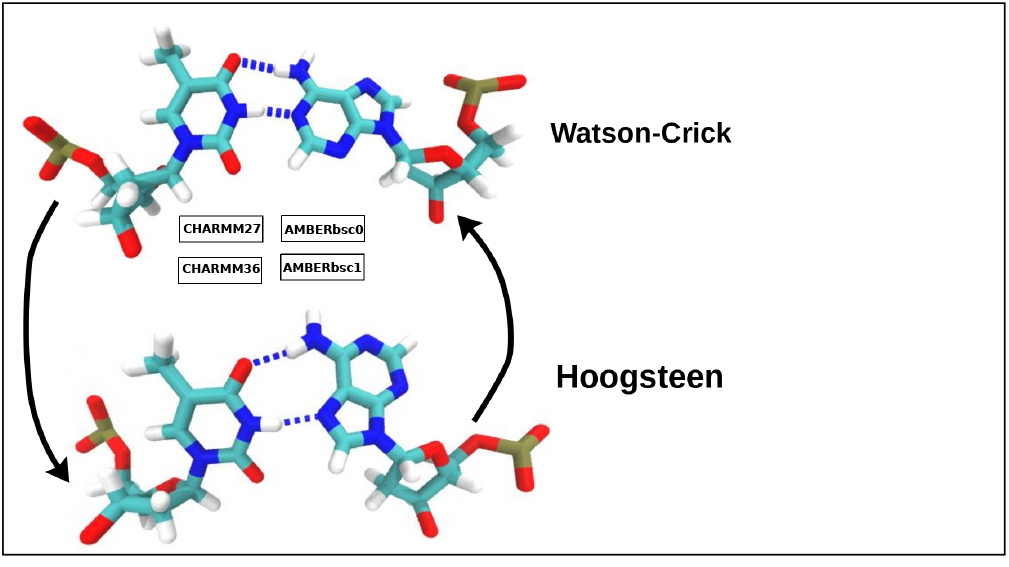

## Introduction

Based on the pioneering X-ray crystallography results of Rosalind Franklin^1^ in 1953, Watson and Crick proposed the structure of duplex DNA,^2^ a model later popularized as the Watson-Crick (WC) structure. By using the bases in their most plausible tautomeric form, they determined complementary base pairs (bp) between adenine and thymine (A−T), as well as guanine and cytosine (G−C).^2^ When the following WC base parings are assembled in a right-handed helix structure, the overall geometry is referred to as B-DNA. Almost 50 years after this discovery, Abrescia et al. have reported a different kind of double-helical structure, from the regulatory regions of DNA, involving a more uncommon Hoogsteen (HG) base pairing conformation.^3^ In relatively recent nuclear magnetic relaxation (NMR) R_1*ρ*_ experiments of free B-DNA, Nikolova et al. showed that both A−T and protonated G−C (G−C^+^) WC base pairs spontaneously formed the HG structure, with a population and lifetime of 0.1-1% and 0.3-1.5 ms, respectively.^4^ Additionally, HG A−T bp are more abundant compared with G−C^+^ bp under solution conditions.

The HG base pairing form was initially observed in isolated nucleotide crystals in the late 1950s using heavy atom X-ray diffraction.^5^ HG structures form when the purine base (adenine or guanine) is flipped over 180 degrees with respect to the glycosidic bond, resulting in a *syn* conformation (as opposed to *anti* in WC) resulting in the shortening of the helix diameter (Figure 1). This structural change creates a chemical environment different from canonical WC bp, and plays a key role in binding to proteins^6,7^ and to small molecule drugs.^8^

**Figure 1:**
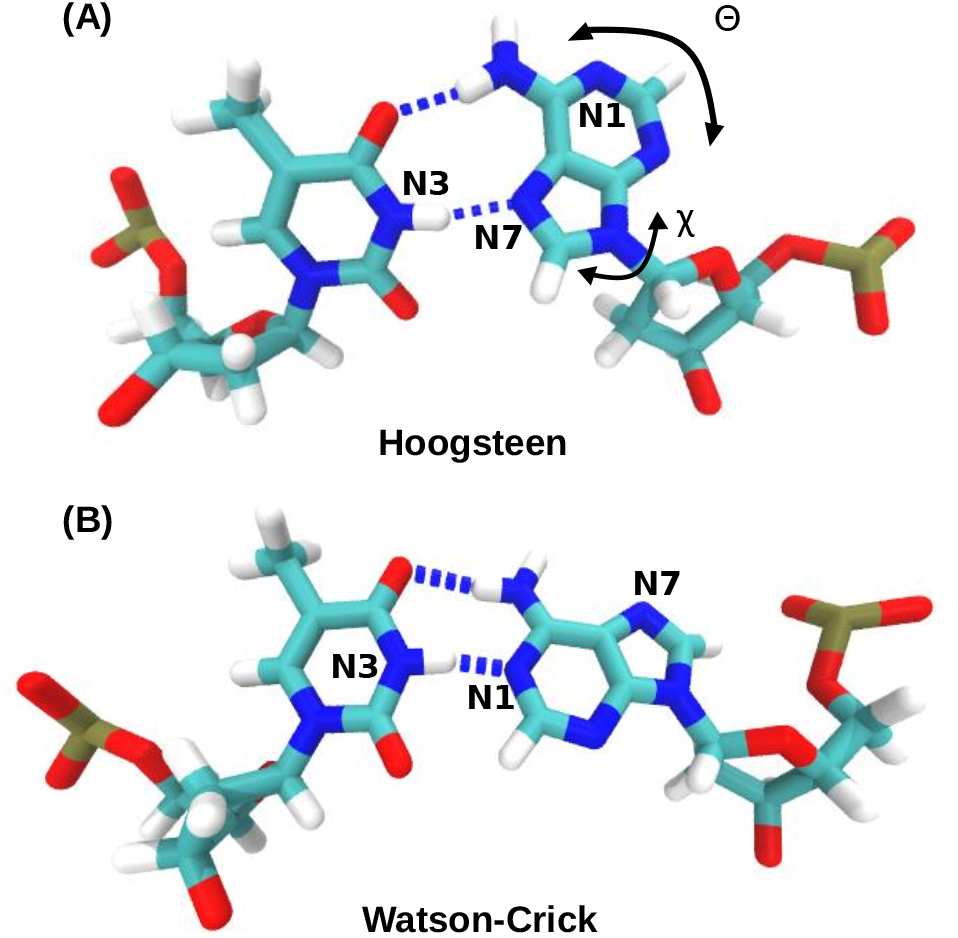
The conformation of (A) Hoogsteen and (B) Watson-Crick A-T base pairs in B-DNA. In the HG bp, the adenine base (on the right side) is flipped over 180 degrees with respect to the glycosidic bond, which results in hydrogen boding with the nitrogen in the five membered ring, and shorter helix diameter. The N1, N3, and N7 nitrogen atoms, referred to in the text, are also marked in the figure.

HG base pairs have also been observed in other scenarios, including damaged DNA,^9–11^ DNA duplex conjugated with the DNA repair machinery,^6,12,13^ and other diverse DNA-protein interactions.^4,6,14–16^ DNA repair processes that do not go up to completion, can lead to cytotoxic damage, potentially resulting in cancer or cell death. An example is the methylation of DNA, which has been shown to cause blocking in DNA replication. Recent work by Xu et al. showed that WC bp are vulnerable to methylation in double-stranded DNA, and the sensitivity to methylation increases for the HG bp. ^7^ Alternatively, a methylated adenine base is more prone to form a HG base pair as a result of stabilizing steric interactions ^17^ Additionally, HG base pairs have been hypothesized to play an important role in DNA repair mechanisms. For instance, the crystal structure of polymerase-*ι* (hPol-*ι*) revealed a replication mechanism dominated by HG base pairs.^18^ The structure of hPol-*ι* encourages replication via the N^2^-adducted G, which forms a syn conformation in DNA, and in turn facilitates a binding mechanism through the minor-groove adducts. ^18^ The HG bp stabilizes the tumor suppressor p53-DNA complex and enhances its binding affinity. ^12^ Binding affinity of small molecules (like intercalation agents and anticancer drugs such as echinomycine) to nucleic acids is also increased in the vicinity of a Hoogsteen base pair.^19^ These examples demonstrate the biochemical significance of the HG configuration in nucleic acids and encourage detailed studies to understand the mechanism of formation of such non-canonical base pairing structures.

Molecular dynamics (MD) simulation can be used to understand the mechanism of biomolecular processes in atomistic detail. The HG base pairing has also been thoroughly investigated using classical MD simulation over the past two decades. The initial set of works by Cubero et al. focused on the structure and stability of the antiparallel helix solely consisting of HG base pairs^20^ and the WC-HG junctions.^21^ The energetics of HG base pairing have also been explored using extensive quantum chemical studies by Hobza and co-workers,^22–25^ leading to the discovery that in gas phase the HG base pair is more stable than the WC conformation.

From experimental NMR relaxation studies, the timescale of the transition between WC and HG base pairing process has been reported to be 50-250 ms, which is beyond the reach of conventional MD simulation with currently available computing facilities. Therefore, the process of transition between these two conformations could only be studied using enhanced sampling methods including umbrella sampling, ^26^ metadynamics^27,28^ and meta-eABF.^29^ Path sampling techniques such as transition path sampling (TPS)^30^ have also been employed to sample the conformational space of the WC to HG bp transition. Yang et al., computed the free energy landscape of the WC to HG transition in A6-DNA, using umbrella sampling and multiple walker well-tempered metadynamics, and captured multiple transition pathways and observed the spontaneous changes in base flipping.^31,32^ Similarly, studies from our group used a fast converging method called meta-eABF^29^ to obtain the potential of mean force (PMF) for the same transition process with CHARMM36 force field parameters.^33^ Our free energy landscape revealed a stable open base pair conformation adjacent to the WC base paired state. The mechanistic analysis using Markov State Modeling ^34,35^ also revealed a melted base pair state to be the primary intermediate in the transition.^33^ This highly stable extra-helical conformation is inconsistent with the conventional idea about the stability of DNA, and provokes further discussion on the accuracy of force fields in describing the base pairing intermediates and transition pathways. Path sampling studies by Vreede et al.^36^ and Hooft et al.^37^ also revealed that partial opening of the A-T base pair is necessary for the conformational switching, but no stable open base paired intermediate conformation was observed.

The force field is an important component of MD simulations, since the accuracy of our results are dependent on the accuracy of the force field model. The force field is the functional forms and the parameter set used to represent the potential energy function of a biomolecule in the context of classical MD simulation^38^. Gradients of this potential energy function are used to propagate dynamical trajectories by solving Newton’s equation of motion.^39^ For nucleic acids, such as DNA and RNA, the commonly used non-polarizable force fields are the CHARMM (CHARMM27, CHARMM36) and AMBER (AMBERbsc0, AMBERbsc1) family models.^40^ Different models have different refinements in order to improve the description of nucleic acid properties. For example, the CHARMM27 model has refined backbone dihedrals, sugar puckering, and glycosidic linkage to match the results of model compounds computed with *ab-initio* calculations.^41^ But it can model the groove width, population of the BI backbone in DNA, and opening of WC bp with limited accuracy. For CHARMM36, improvements were focused on refining backbone dihedral angles, sugar puckering, and the 2’-hydroxyl dihedral for RNA parameters. ^40^ In the AMBER family, AMBERbsc0 was focused on refining the backbone conformations (*α* and *γ* dihedrals), in order to lower overpopulated *α* gauche+ and *γ* trans conformations. ^42,43^ Additionally, the AMBERbsc1 model was constructed with AMBERbsc0, but with refined sugar puckering and *χ, E*, and *ζ* dihedrals.^44^ These refinements have become an essential process in order to obtain accurate results in the microsecond timescale. Comparative studies of the common nucleic acid force fields have been performed to evaluate the mesoscale properties of DNA,^45^ and to reproduce NMR structures of the Drew-Dickerson dodecamer (DDD) system.^43^ Minhas et al. found the CHARMM27 model to have the most stable trajectories and best agreement to the experimental data for describing the mesoscale properties of multimicrosecond simulations of long DNA fragments (40 base pairs) in a cubic solvation box of TIP3P water.^45^ Additionally, the CHARMM27 model had the best agreement with X-ray diffraction and NMR results compared with CHARMM37, AMBERbsc0, and AMBERbsc1 models. While the study revealed both AMBER family models to have good agreement with the experimental data, the CHARMM36 model had unstable trajectories in the microsecond timescale and produced irreversible fluctuations.^45^

Whether the choice of force field can affect the pathways and free energy landscape of the conformational transition between the WC and HG base pairing forms is not yet clear. But such understanding is vital, considering the important physiological role of the HG base paring in DNA repair and replication. In the current work, we address this question by comparing the underlying free energy landscape of WC to HG transition, and the energetic and entropic stabilization of individual base pairing states for the four different force fields: CHARMM27, CHARMM36, AMBERbsc0 and AMBERbsc1. Although newer nucleic acid force fields are available, ^43,46^ our choice of force fields was inspired by the recent comparative study by Minhas et al., ^45^ and molecular dynamics studies of HG base pair formation in the literature. The free energy surface and associated thermodynamic properties of the HG conformation of the A16−T9 base pair in A_6_-DNA^4^ are explored through enhanced sampling simulation using the advanced meta-eABF approach. The potentials of mean force (PMF) obtained from the meta-eABF simulations are used to compare the force field models in terms of the minimum free energy pathway for the WC-HG transition. Presence of HG and WC base pairs is observed in all the four models, although the relative stability of each base pairing configuration can be somewhat different. To obtain a fundamental rationale behind this discrepancy, we computed the quasi-harmonic entropy and the root-mean-square fluctuations (RMSF) of the two base pairing conformations from equilibrium trajectories for all force field combinations. We observe that indeed there is a large variation of the free energy, enthalpy, and entropy of HG formation among the different force fields, although the predicted structures of the HG and WC conformation are almost identical. Based on our findings, we explain the main distinctions between each force fields in describing the molecular details of WC and HG base pairing conformations and compare our findings with experimental results.

## Computational Methods

### System Preparation and Equilibration

We performed our simulations on the A6-DNA fragment (PDB ID: 5UZF)^47^ reported in the NMR relaxation study by Nikolova et al.^4^ This structure includes six A-T base pairs in a 12 bp sequence (CGATTTTTTGGC). The topology files for the CHARMM and AMBER family force fields were prepared using Visual Molecular Dynamics (VMD) ^48^ and AmberTools (tleap)^49^ respectively. The initial structure of A6-DNA was placed in a solvated box with TIP3P water and 22 Na+ ions to neutralize the overall system. CHARMM-TIP3P force field^50^ is used to model the water and ions when the CHARMM family force field is used for nucleic acids, and the TIP3P model^51^ is used for water and ions in the simulations with AMBER force field. We did not add additional salt in the system, as previous computational studies under neutralizing ion concentration could most accurately reproduce the experimental population ratio between WC and HG conformations.^32^ Also, the NMR experiments were performed at 25 mM ionic strength, which should correspond to ¡10 ions in our simulation box while we already have more than 20 neutralizing Na+ ions. A water padding of 17Å was included in each direction. Each solvated system comprised of ~26000 atoms. In case of the CHARMM family force fields, a 12 Å cutoff was used for non-bonded forces along with a switching function with switch distance of 10 Å. For AMBER family force fields, a 9 Å cutoff was used for non-bonded forces without any swithing function, following the recommended protocol.^52^ The structures were minimized for 50,000 steps using the conjugate gradient algorithm, followed by a slow heating to reach 298K temperature using a velocity rescaling thermostat with periodically updating the temperature at a rate of 1 K/ps. During the slow heating harmonic restraints of 3 kcal mol^*−*1^ Å^*−*2^ were applied on the heavy atoms. These restraints were then removed over a period of 1.2 ns, at a rate of 0.5 kcal mol^*−*1^ per 200 ps. Then, the system was placed in a NVE Ensemble for 3 ns with harmonic restraint with force constant of 0.1 kcal mol^*−*1^Å^*−*2^, followed by an unrestrained NPT simulation for 10 ns. The terminal base pairs were restrained with a force constant of 0.05 kcal mol^*−*1^Å^*−*2^ for all equilibration, production and enhanced sampling simulations. The temperature was kept constant at 298 K using the Langvein thermostat with a coupling constant 1 ps^*−*1^. The pressure was controlled to be at 1 atm, using the Nosè-Hoover Langvein piston with piston decay of 50 fs and oscillation period of 100 fs.^53,54^ Identical protocol was used for all four force field models. All molecular dynamics simulations reported in this study have been performed using the NAMD 2.14 software package.^55^ The input files for all simulations are provided with the manuscript (See Data Availability Statement).

### Enhanced Sampling Simulation

To accelerate the rare transitions between the WC and HG form, we performed enhanced sampling simulations using a combination of metadynamics ^27^ and extended-system adaptive biasing force (eABF),^56^ commonly known as the meta-eABF approach.^29^ Meat-eABF simulations were performed using the colvars module ^57^ patched with the in NAMD 2.14 simulation package. The 2-dimensional potential mean of force (PMF) was calculated along the following collective variables (CV): the glycosidic angle *χ* and a pseudo-dihedral angle Θ of the A16-T9 base pair. ^31^ The Θ angle characterizes the base flipping into the major and minor grooves of DNA, while the *χ* dihedral angle characterizes the rotation of the adenine base with respect to the deoxyribose sugar (O4’,C1’,N9, and C4). A pictorial representation of these two torsion angles are provided in the following reference 32. We started the meta-eABF simulations from the end points of NPT equilibration and propagated for ~200-400 ns, until the PMFs were sufficiently converged. Gaussian hills of height 0.06 kcal/mol and width 3 CV unit (15°) were deposited every 2 ps for the metadynamics part. Additionally, the ABF bias was applied against the average force after 1000 samples were collected for the corresponding bin of width 5°. The MEPSA program^58^ was used to obtain the minimum free energy pathway from the 2D energy landscape obtained from enhanced sampling simulation. The pathway is predicted as a smooth curve joining various nodes on the free energy surface using the “node-by-node” search algorithm,^58^ which bears similarity with the Dijkstra algorithm.^59^ The start and the end point of the interpolated pathway is manually chosen to be, respectively, the minimum of the WC basin and the minimum energy point approximately at *χ* = 180°. This is done to avoid any spurious interpolations resulting from the inability of the algorithm to take into consideration the periodicity of the collective variable.

### Equilibrium Simulations for Entropy and Energy Calculation

Structures of the WC and HG base paring conformations were sampled from the meta-eABF trajectories based on the hydrogen bond donor acceptor distances. Total 8 structures for four different force fields were used as starting points for 100 ns unbiased MD simulations with identical simulation setup as the meta-eABF simulation. The coordinates were saved every 10 ps. The hydrogen bonding donor distances between atoms N1(A)-N3(T) and N7(A)-N3(T), were monitored to ensure WC and HG conformations remained intact throughout the simulation (see Supporting Information). These trajectories were used to calculate the root-mean-square fluctuation (RMSF) and quasi-harmonic entropy. ^60^ Five 50 ns trajectory segments were extracted from the last 90 ns of each trajectory using overlapping time windows: 10 ns - 60 ns, 20 ns - 70 ns, 30 ns - 80 ns, 40 ns - 90 ns, 50 ns - 100 ns. The entropy and RMSF were computed from each segment, and the mean and 95% confidence intervals are reported for the 5 data points.

The RMSF of all heavy atoms were computed using GROMACS 2018.1 software^61^ with the `gmx rmsf` tool. Details of the calculation of quasiharmonic entropy from the covariance matrix of atomic fluctuation is provided in the original paper.^60^ In brief, the entropy is given by,

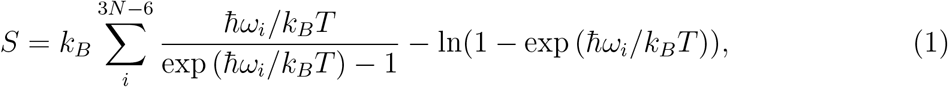

where *k*_*B*_ is Boltzmann constant, *ħ* is *h/*2*π* where *h* is Planck’s constant and *T* is absolute temperature. The frequencies *ω*_*i*_ are obtained as 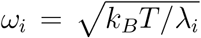, for *i* = 1, 2, 3, …, 3*N* − 6 where *λ*_*i*_ are eigenvalues of the mass weighted covariance matrix *σ* obtained from MD simulation. The matrix *σ* is given by **M**^1*/*2^*σ*′**M**^1*/*2^ where **M** is the mass-matrix (diagonal matrix containing the masses of the different atoms) and *σ*′ is the covariance matrix of atomic coordinates. The elements of *σ*′ are given by

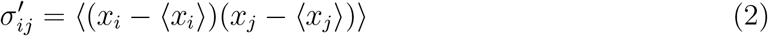

*x*_*i*_ is the *i*th component of the 3N dimensional Cartesian coordinate space of N atoms and ⟨⟩ indicates ensemble average. The number of atoms involved in the calculation (*N*) equals to the number of heavy atoms in the DNA. The entropy is dominated by slow vibrational modes, so we excluded the fast moving hydrogen degrees of freedom from our analysis. The covariance matrices and their eigenvalues were computed using the `gmx covar` module of GROMACS 2018.1 package.^61^ The entropy was then evaluated using equation 1 by an in house python code. Non-bonded interaction energies between bases and nucleotides were calculated using the NAMDEnergy module of VMD.^62^

## Results and Discussions

### Free Energy Surfaces

Similar to our previous study, ^33^ the meta-eABF approach could produce the PMFs within a few hundreds of nanoseconds, at significantly less computational cost compared to other approaches,^31,32^ facilitating the comparative study between multiple force fields. The free energy surfaces obtained for different models are shown in Figure 2. In all models, the Watson-Crick (WC) base paired configurations were present near Θ ~ 0° and *χ* ~ −100°, and the Hoogsteen (HG) form near Θ ~ 0° and *χ* ~ 60°. The free energy difference between the WC and HG base pair (ΔG_*WC→HG*_) was determined to be ~ 8 kcal/mol for CHARMM27, ~ 6 kcal/mol for CHARMM36, ~ 1 kcal/mol for AMBERbsc1, and ~ 0 kcal/mol for AMBERbsc0. When comparing the NMR value, 3.3 kcal/mol, the free energy difference is overestimated in the CHARMM family FFs and underestimated in both AMBER family models, and closest in agreement to the CHARMM36 and AMBERbsc1 force field. These discrepancies are much larger compared to the results obtained by Yang et al.^31,32^ using more expensive umbrella sampling ^26^ and multiple-walker well tempered metadynamics^28^ simulations and can possibly indicate an artifact of the meta-eABF approach which converges faster but only provides a qualitatively accurate free energy landscape for systems with high energy barrier.^63^ Also, the uncertainty of the free energy landscapes, we computed, is at least ~ 1 kcal/mol as apparent from the convergence plots (See Supporting information).

**Figure 2:**
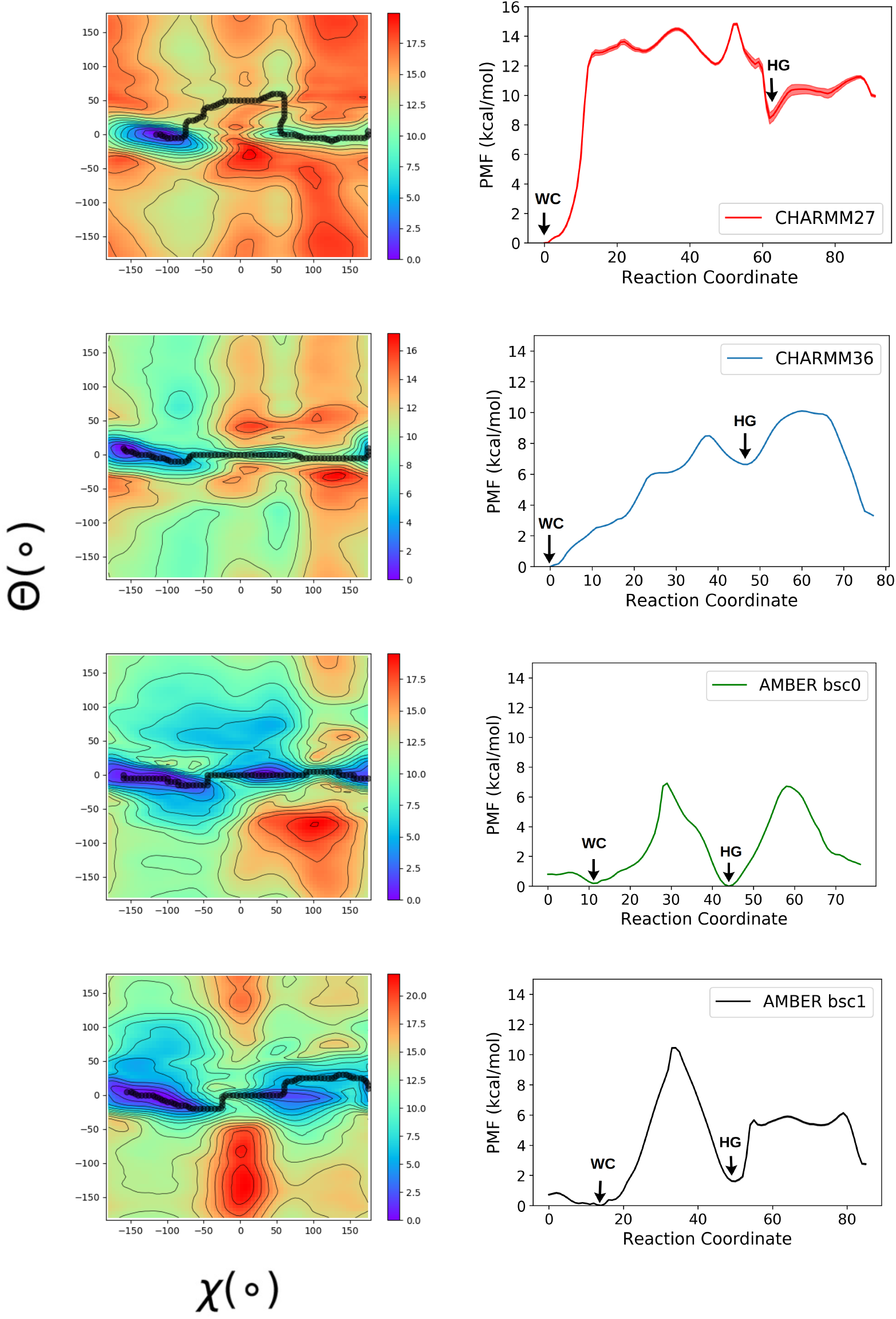
The free energy surface of the A16-T9 base pair in A6-DNA was obtained using meta-eABF method for different force field models. The error bars obtained from the last 50 ns of the simulation using MEPSA software, are indicated in the 1D plots. But they are too small to distinguish from the plot, except for CHARMM27 force field.

All the four free energy landscapes indicate that the opening of the base pairs through the major groove region is more energetically favorable compared to the minor groove region, especially when starting from a WC structure, which is the most prevalent form in solvated DNA duplex. This pathway is in agreement with the previous experimental^4^ and simulation studies.^33,36^ Except for CHARMM36, all three force fields show that the base paired structures (WC or HG) are more stable than the open base paired configurations. In CHARMM36 we can observe an open base pair state in the major groove region near the WC form which is lower in free energy in comparison to the HG state. This result is consistent with the previous work, where a very stable open base paired intermediate state between WC and HG was captured using CHARMM36 force field and Markov State Modeling. ^33^

Additionally, for the majority of models, the minimum free energy pathway reflected in the transition mechanism is a direct 180° flip-over of the adenine glycosidic angle whereas the CHARMM27 FF shows the minimum free energy pathway involves slight opening of the base pair indicating a possible extra-helical mechanism. From previous analysis of the T9-A4’ base pair in sequence 5’-CGATTTTTTGGC-3’, it was concluded that the extra-helical mechanism, where the purine base flips out of the DNA helix first, was preferred for the HG to WC base pair transition.^36^ Although this mechanism was not the most energetically favorable in CHARMM36, AMBERbsc0, and AMBERbsc1, the metastable open base pair conformations observed in those FFs allows for the possibility of conformational switching between WC and HG form via an extra-helical intermediate. For all the force fields, shallow free energy regions are detectable in the major groove (Θ *>* 0) and minor grooves (Θ *<* 0). In the CHARMM36 and AMBERbsc1 force fields, the open base pair state, adjacent to the Waston-Crick minima, is clearly distinguishable.

In our previous study,^33^ we suggested that the dihedral angle based reaction coordinates may not be able to uniquely distinguish between WC and HG states as some structures which do have the necessary hydrogen bonds for base pair formation can also be assigned to either of these two categories just from the *χ* and *θ* torsion angle values. Here, we propose to use the hydrogen bond donor acceptor distances to identify the base paired forms. In WC base pairing, an N-H-N type hydrogen bond forms between the N1 nitrogen of adenine and the N3 nitrogen of thymine nucleotides (N1(A)-N3(T)). In the HG form, the N3(T) forms a hydrogen bond with N7(A) instead. So, by measuring the distance between the N1(A)-N3(T) atom pair and N7(A)-N3(T) atom pair the WC and HG states can be uniquely identified. This is reflected in the time series plots of dihedral angles and hydrogen bond donor acceptor distances in Figure 3. We projected our free energy landscapes from Figure 2 in the hydrogen bond donor acceptor distance space (Figure 4). All four models clearly distinguish the WC and HG states as free energy minima in the configurational space. The relative free energies of the HG base pairs are in slightly better agreement with the experimental results (CHARMM27: ~6 kcal/mol, AMBERbsc1: ~2 kcal/mol, AMBERbsc0: ~1 kcal/mol, CHARMM36: ~6 kcal/mol), reinforcing our proposition that in *χ*-*θ* based representation the WC and HG states get contaminated by other states with higher/lower free energy and provides a poorer estimate of the relative free energy. One should note that these distance based coordinates may not be the optimal choice as collective variable for the application of the biasing force, as we indeed need to force the transition of the glycosidic torsion angle to go from one base pairing conformation to the other. The hydrogen-bond donor-acceptor distance based CVs are useful to distinguish between WC and HG conformations during the post-processing of the MD trajectories. In this sense, the primary significance of Figure 4 is that it shows the free energy landscape of the process of switching between WC and HG base paring along the H-bond donor acceptor distance collective variables which better distinguish the the two basins as opposed to traditionally used torsion angle based CV. It was necessary to reweight the meta-eABF trajectories biased along *χ* − *θ* coordinates, as biasing these distances directly may disrupt the base paired structure of the DNA duplex making it difficult to revisit the base paired states during the simulation. This leads to significant noise in the free energy landscape in Fig. 3 as the 2D space in the distance based CVs is not uniformly explored. Nevertheless, the energy values we observe specifically at the WC and the HG minima are slightly more reliable than what we observe in Fig 2 because of the better ability of these CVs to distinguish these states.

**Figure 3:**
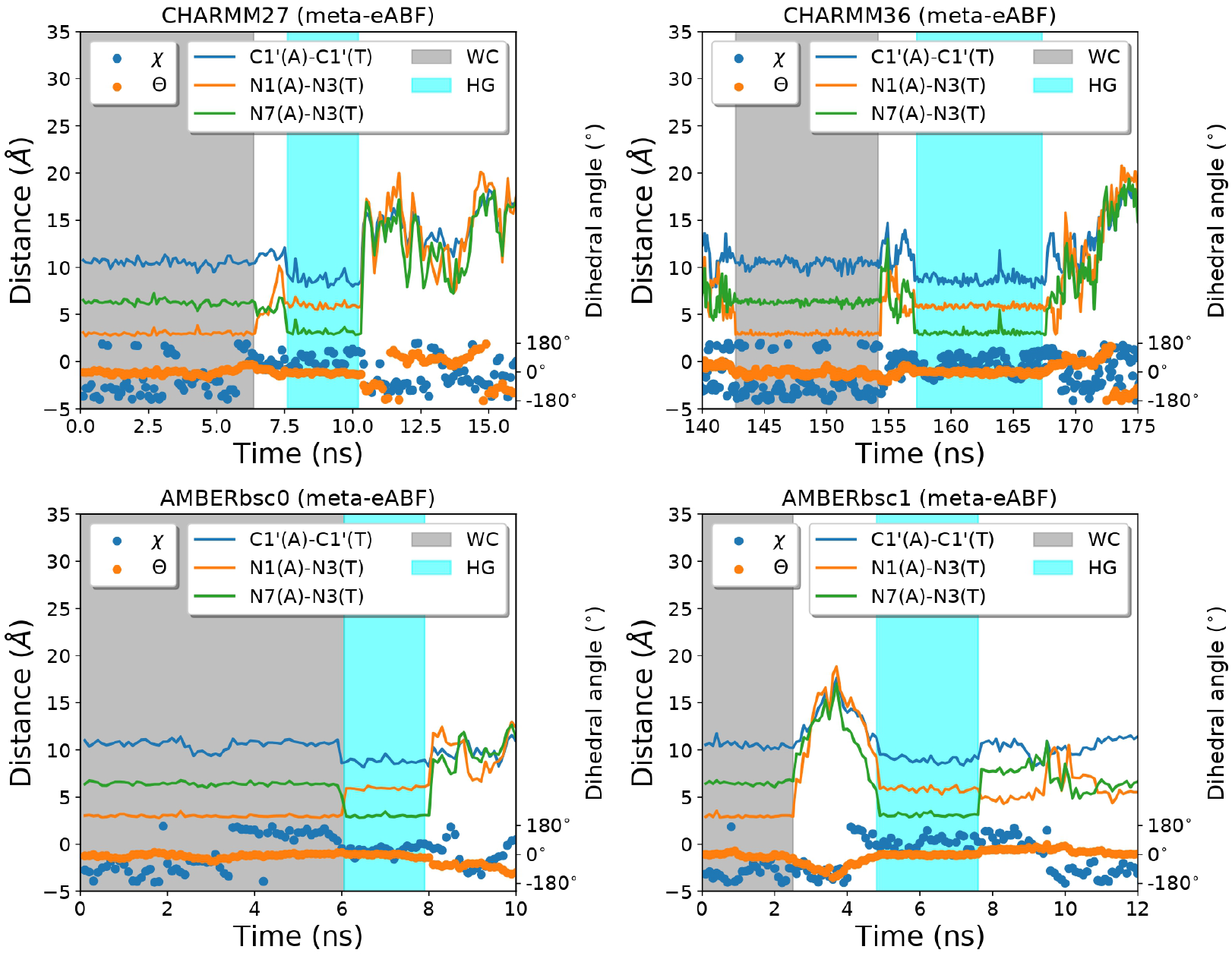
Representative regions of the meta-eABF trajectories showing transitions between WC and HG conformations. The hydrogen bond donor acceptor distance, the helix diameter measured as the C1’-C1’ distance, and the two torsion angles *χ* and *θ* are depicted as a function of time.

**Figure 4:**
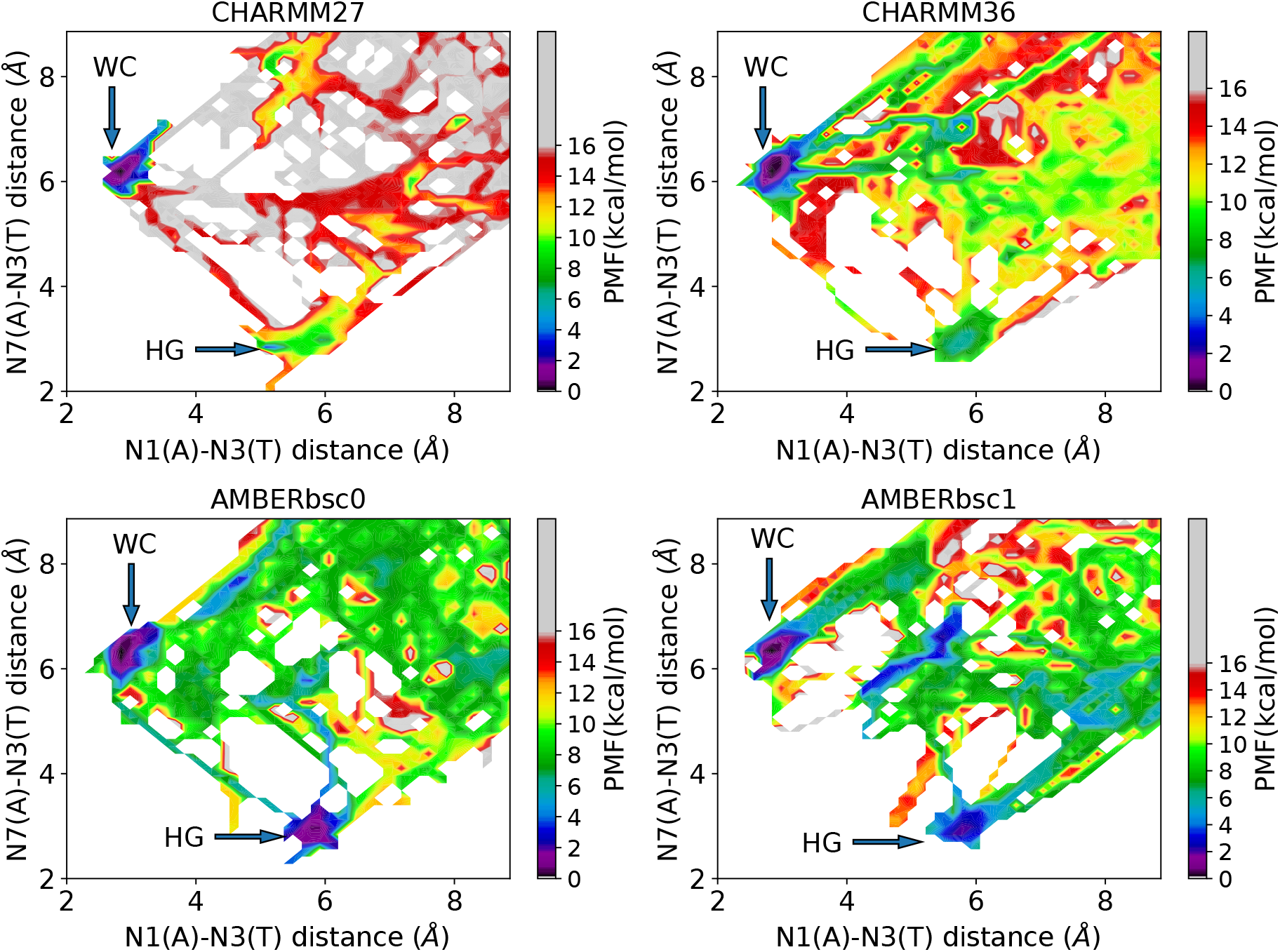
The free energy landscape of the conformational switching between WC and HG base pair configurations, projected onto the two hydrogen bond donor acceptor distances, for different force field models.

### Root Mean Square Fluctuations and Absolute Entropy

The RMSF of heavy atoms are different at the A16 base and the nearby base pairs when comparing both WC and HG conformations, Figure 5. For instance, in both the CHARMM family force fields, the A16 base experienced more fluctuations in the HG form than the WC conformation. The opposite pattern was observed in the AMBER family models, where the fluctuations at the A16 base were larger in the WC form. In all force fields, there are also fluctuations occurring near the A16 and T9 base, which indicate the impact of the Hoogsteen form on the entropy of the B-DNA. Large change in fluctuations could occasionally be observed in individual atoms in neighboring bases, for example, in the A18 base in case of AMBERbsc0 force field (Fig. 5). Such fluctuations can be a result of long-range and short range correlated motions of the atoms, but it is difficult to pinpoint from the current results. Similar collective motion between a base flipping out, and the neighboring bases were observed by Lavery and coworkers in their “Saloon door” mechanism, although we do not observe any base flip out motion in our unbiased trajectories. It is more interesting to look at the increase/decrease of fluctuations at an atomic resolution. The change in fluctuation depicted in Fig 5 in residue-specific information, we show it in Fig. 6 as an atom-specific scale to understand which exact atoms are gaining and losing fluctuations. One common theme is apparent for all force fields: The base regions (particularly the six membered ring of the adenine base) becomes more flexible in the HG form than in the WC form, while the backbone region becomes slightly more rigid in the HG form. The effect is not very consistent for the Thymine base (Figure 6). This indicates that the apparent rigidity of the HG base pair is not equally contributed by all the atoms.

**Figure 5:**
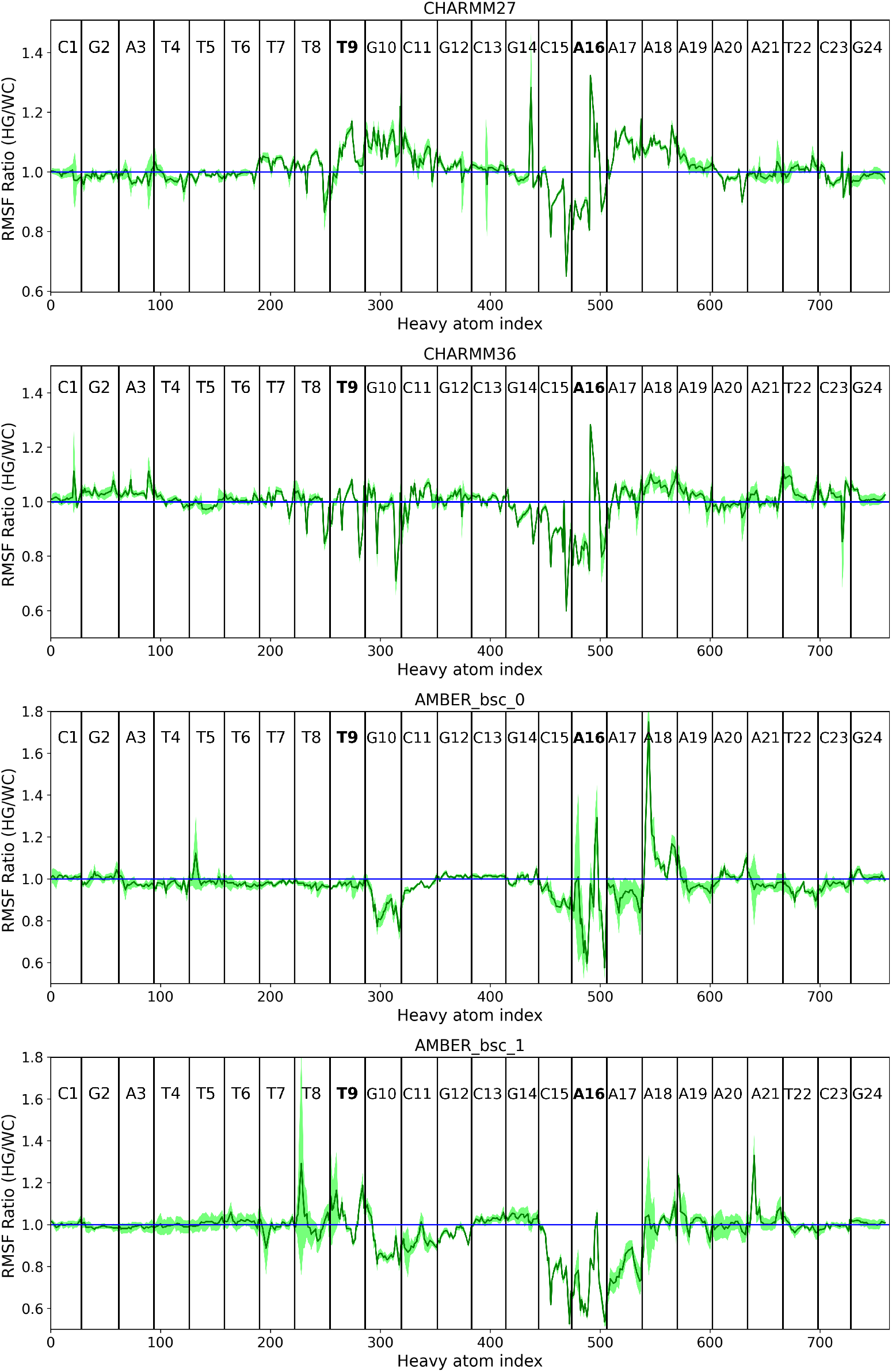
The RMSF ratio of the DNA duplexes involving the HG to WC form of the A16-T9 base pair. The location of all the base pairs are indicated by vertical lines. The A16 and the T9 nucleotides have been labelled in bold font. The error bar computed as the 95% confidence interval of the five 50 ns trajectory segments is indicated as the light green shading.

**Figure 6:**
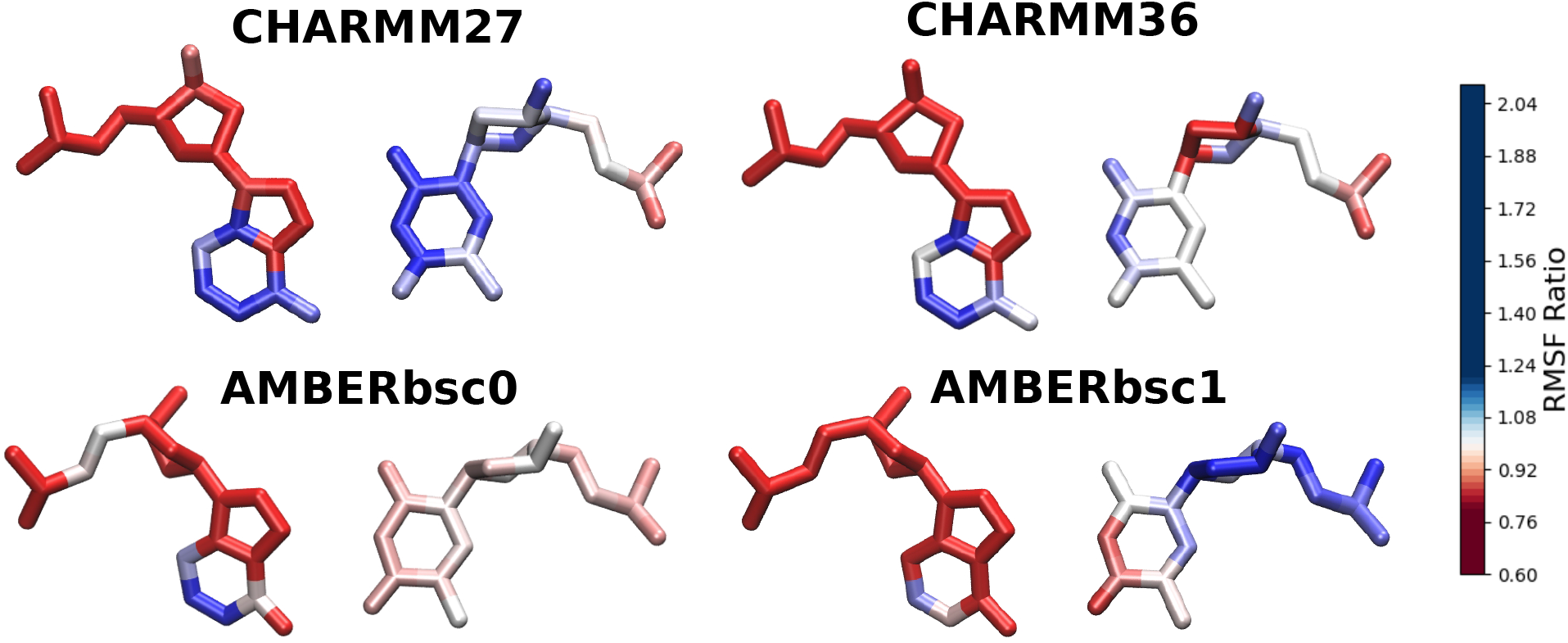
The atoms of the A16-T9 base pair colored according to the RMSF ratio between the WC and HG state. The blue color indicate RMSF ratio greater than 1 and red color indicate less than 1.

In Figure 7, the quasi-harmonic entropy of the DNA duplex involving one Hoosgteen conformation, is found to be lower than that of a full WC DNA duplex, for all four force fields, possibly due to the increase of rigidity caused by the shrinkage of the helix diameter at the HG state. The extent by which the HG base pairing form is entropically destabilized varied between different force field models. The entropy difference of HG and WC base pairs is lowest in the CHARMM27 ff and is highest in AMBERbsc1, in contrast to the free energy results that indicate a lower free energy difference in AMBER ffs compared to CHARMM ffs. To understand the role of entropy and enthalpy in the stabilization of the WC base pairing form over the HG, we performed non-bonded interaction energy calculations. We found that the interaction energy between two bases are stronger for the HG base pairing form than the WC base pairing form, for all force field models (Fig. **??**). Although this seems counterintuitive, previous studies of Gould and Kollman show that in the crystalline state the HG base pairing form is energetically more stable than the WC form. ^64^ This stabilization is much higher for the AMBER family force fields than the CHARMM family force fields. When we look at the interaction energy of the base pair with the rest of the DNA duplex, the CHARMM family force fields show a positive (destabilizing) change in the interaction energy going from WC to HG, while a negative (stabilizing) change is observed for the AMBER family ffs. We also computed the relative “stacking” energy between the two base pairing forms. Although it is necessary to employ high level quantum mechanical approaches to calculate the accurate *π* stacking energy of a given base pair, ^22,65,66^ we are limited to the use of nonpolarizable classical force fields for the current study. We therefore use the non-bonding interaction energy between the conjugated aromatic rings between the A16-T9 base pair and the two adjacent base pairs (i.e. A17-T8 and C15-G10). We refer to this as the “base stacking” energy in the rest of the manuscript, to differentiate from the “*π* stacking” energy. The difference of base stacking energy between the WC and HG form in CHARMM family FFs is almost negligible. But the stacking energy for the AMBER family FFs is stronger (more negative) in the HG form compared to the WC form (Fig. **??**). These results clearly suggests that the two types of force fields treat the HG base pairing very differently in terms of energy costs. The positive (or less negative) energy cost of forming a HG base pair in CHARMM FFs results in a higher free energy difference between the WC and HG base pairs despite a lower entropic cost compared to AMBER FFs.

**Figure 7:**
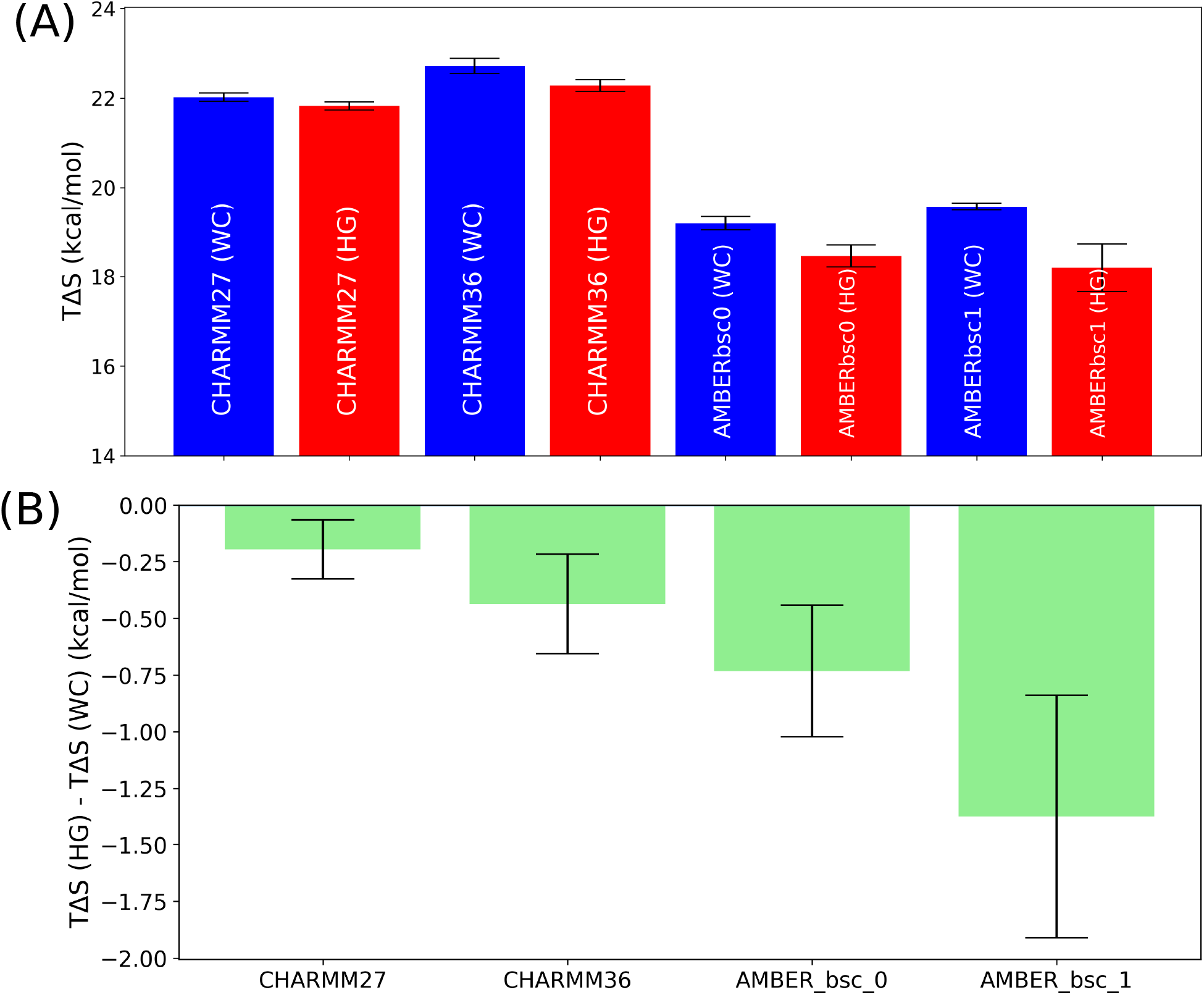
(A) The absolute value and (B) The relative value of the quasi-harmonic entropy of the HG base pair with respect to the WC base pair for different force fields.

**Figure 8:**
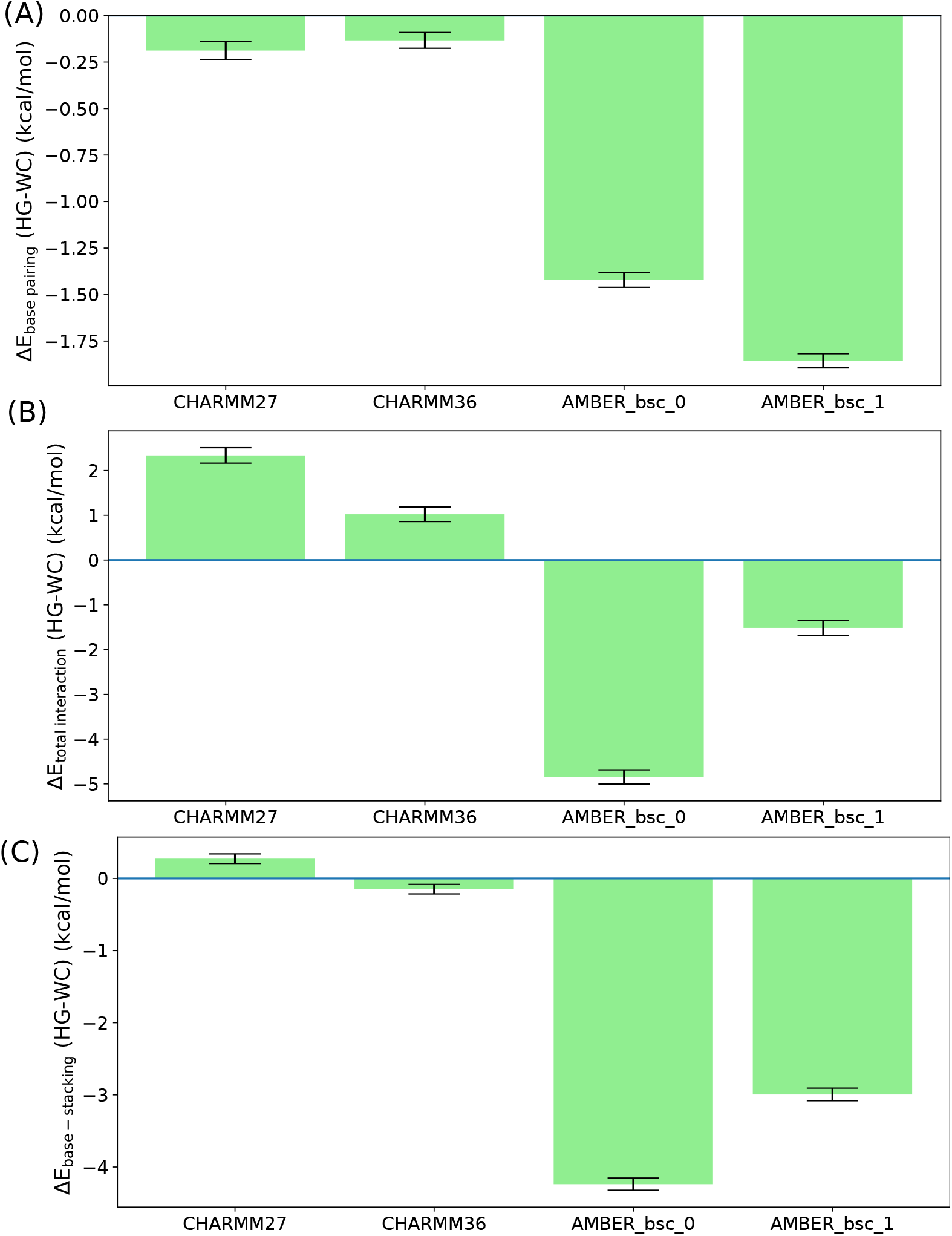
The non-bonded interaction energies in the HG base pair relative to the WC base pair, for the following interactions: (A) between the A16 and T9 bases, (B) between the base pair with the rest of the DNA duplex, and (C) the stacking energy defined as the interaction energy between the *π* conjugated region of the base pair with the two adjacent base pairs.

Nevertheless, the structural properties of the WC and HG base pairs are fairly consistent throughout the four force field models. The two torsion angles and various inter-atomic distances between the A16 and T9 bases are reported in Table 1 and 2, respectively. Except for a wider fluctuation and a shifted location of the HG minima in the *χ* − *θ* space for the AMBERbsc0 force field, the results of different force fields are in agreement with each other. One should also consider that the results in Table 1 and 2 were averaged over multiple frames of a single MD trajectory. Due to the correlated nature of the data points, the true uncertainty is likely higher than the reported standard deviation, indicating that any apparent disagreement of structural parameters between force fields is insignificant. The time evolution of the H-bond donor acceptor distances, the helix diameter (measures as the distance between the C1’ atoms), and the torsion angles for the 100 ns equilibrium trajectories are provided in the supporting information. It is clear from these plots that no WC to HG transition or base opening takes place during the simulation, confirming that these structures indeed represent minima in the free energy landscape.

**Table 1:** The values of the glycosidic angle *χ* and the base flip-out angle *θ* for the WC and HG conformation for the four different force fields. The results are reported as mean ± standard deviation of the structures sampled in the 100 ns unbiased MD simulation.

| Force Field | WC<br>$\chi(^{\circ})$ | HG<br>$\chi(^{\circ})$ | WC<br>$\theta(^{\circ})$ | HG<br>$\theta(^{\circ})$ |
| --- | --- | --- | --- | --- |
| CHARMM27 | -109.2±13.1 | 46.7±10.3 | -2.2±3.9 | 0.0±2.4 |
| CHARMM36 | -113.0±14.8 | 45.2±10.7 | -0.4±3.8 | -0.2±2.3 |
| AMBERbsc0 | -113.3±20.1 | 26.8±22.8 | -5.9±5.2 | -0.3±2.2 |
| AMBERbsc1 | -110.3±15.7 | 61.0±9.3 | -5.3±6.0 | -1.3±2.0 |

**Table 2:** The distances between the hydrogen bond donor-acceptors (N1-N3 and N1-N7), and the C1’ atoms of the A16 and T9 nucleotide, for the WC and HG conformation for the four different force fields. The results are reported as mean ± standard deviation of the structures sampled in the 100 ns unbiased MD simulation.

| Force Field | WC<br>N1(A)-N3(T)<br>($\text{\AA}$) | HG<br>N1(A)-N3(T)<br>($\text{\AA}$) | WC<br>N7(A)-N3(T)<br>($\text{\AA}$) | HG<br>N7(A)-N3(T)<br>($\text{\AA}$) | WC<br>C1(A)-C1(A)<br>($\text{\AA}$) | HG<br>C1(A)-C1(A)<br>($\text{\AA}$) |
| --- | --- | --- | --- | --- | --- | --- |
| CHARMM27 | 2.96±0.13 | 5.90±0.25 | 6.31±0.15 | 3.12±0.30 | 10.60±0.28 | 8.88±0.51 |
| CHARMM36 | 2.96±0.13 | 5.87±0.21 | 6.29±0.16 | 3.08±0.22 | 10.60±0.29 | 8.85±0.40 |
| AMBERbsc0 | 2.96±0.12 | 5.84±0.17 | 6.48±0.15 | 3.02±0.16 | 10.56±0.29 | 9.03±0.31 |
| AMBERbsc1 | 2.97±0.12 | 5.89±0.17 | 6.51±0.14 | 3.05±0.14 | 10.59±0.29 | 9.04±0.30 |

## Conclusions

We performed enhanced and regular sampling equilibrium molecular dynamics simulations to understand the role of force field model in predicting the properties related to the formation of the Hoogsteen base pairing in the A6-DNA duplex. Although the equilibrium structures of the HG and WC base pairing forms were consistent in the different force field models, the relative free energy, entropy and interaction energy of the HG base pair change drastically when a different force field model was used. The relative stability of the two base pairing forms are determined by a fine balance of the entropy cost of forming the structurally rigid HG base pair and the gain in interaction enthalpy over the WC structure. The topology of the underlying free energy landscape of the conformational transition between WC and HG when DNA breathes is more or less consistent between the different force field models. However, the predicted free energy costs and the barriers varied widely. Any quantitative comparison at the level of free energy should be performed with caution, given the approximate nature of the PMFs obtained from meta-eABF simulation. Multi-microsecond well-tempered metadynamics simulations can produce more accurate free energy landscapes, but can become prohibitively expensive when trying to compare the results between different force fields. Importantly, the collective variables used for biasing are not optimal for distinguishing the two conformations and more sophisticated artificial intelligence driven order parameters, such as those obtained using spectral gap optimization^67^ and harmonic linear discriminant analysis^68^ may provide a better free energy difference.

It should also be taken into consideration that ergodicity, on the theoretical level, can be proven only as a necessary condition, but never a sufficient one. This means that faster convergence in any simulations does not always guarantee that the “converged” result is the ensemble average, which is, presumably, the experimental result. However, the experimental result is, at the same time, prone to its statistical inaccuracy. Therefore, despite the fact that our results do not agree perfectly with the previous computational studies with the AMBER bsc1 force field (which till now produced better agreement between simulation and experimental free energy difference), the question still remains on the match between MD and bulk experiments. Given this understanding, we here merely demonstrate that the choice of the force field can impact the results of the free energy landscape of DNA of breathing dynamics. When an identical protocol is used, the free energy landscapes from different force fields are comparable, provided that they visit all relevant configurations in the process (see the ergodicity argument above), which we can capture from our free energy landscapes. We chose meta-eABF because it quickly converges to a free energy surface that allows us to make a qualitative comparison between the different force fields in terms of their ability to describe the process in question. Alongside, we expect that both CHARMM and AMBER family force fields lead to a similar degree of exploration of the configurational space when subjected to identical enhanced sampling protocols.

Apart from the relative populations of the WC and HG base paired conformations, the transition kinetics between these two state have also been obtained from NMR relaxation experiments^4^ (exchange rates between 4 and 20 s^*−*1^). But we restricted our analysis only to the thermodynamic properties, as calculating accurate kinetics for a multi-millisecond process with different force fields is computationally extremely expensive. Nevertheless, in our previous work, we employed Markov State Modeling to compute the kinetics of this conformational switching within two orders of magnitude agreement with the experimental data, from the CHARMM36 force field.^33^

Based on the findings of the current study, we could only comment on the fact that the relative stability of the WC and HG base pairing conformation can be affected by the choice of the force fields. But, due to the quality of the agreement with experimental populations being poor in all force fields, it remains difficult to conclusively suggest which force field optimally describes this very important conformational switching in DNA base pairs. Our results rather highlight the diversity of the predictions of molecular properties of WC and HG base pairs from different force fields, and facilitates the assessment of the reliability of future computer simulation studies on DNA breathing dynamics.

### Special Issue Note

The authors declare that they are committed to including, supporting and advancing women in Chemistry and scientific research in general. The authors are aware of the multitude of challenges faced by women in progressing their career in scientific research, particularly in academia, with only a small fraction of the leadership positions held by women. The lead author of this paper, SES, is a first generation female college student from a historically underrepresented community, and has first-hand experience of the enormous difficulties involved. The research group of IA have hosted many female scientists who proceeded to become successful independent scientists. The authors are not complacent, but aware that much more is needed to be done. All authors pledge to undertake conscious efforts in removing the barriers and create an inclusive and supportive environment for women in science.

## Acknowledgement

This work was supported partially by the National Science Foundation (NSF) via grant MCB 2028443 awarded to IA. SES acknowledges support through Pfizer - La Jolla Academic Industrial Relations (AIR) Diversity Research Fellowship in Chemistry for Undergraduates. The authors thank the University of California Irvine High Performance Computing (HPC) facility and the Triton Shared Computing Cluster (TSCC) in the San Diego Supercomputer Center (SDSC) for providing the computational resources. The authors declare no competing financial interest.

## Data and Software Availability

The open-source package NAMD 2.14^69^ is used for all molecular dynamics simulations. The simulation input files and analysis scripts are available from the github repository: https://github.com/dhimanray/DNA-HG-FF.git.

## Supporting Information Available

See supporting information for additional plots of the stability of the HG and WC structures, convergence of entropy calculations, and the absolute values of interaction energies.

