## Supporting information for "Force Field Dependent DNA Breathing Dynamics: A Case Study of Hoogsteen Base Pairing in A6-DNA"

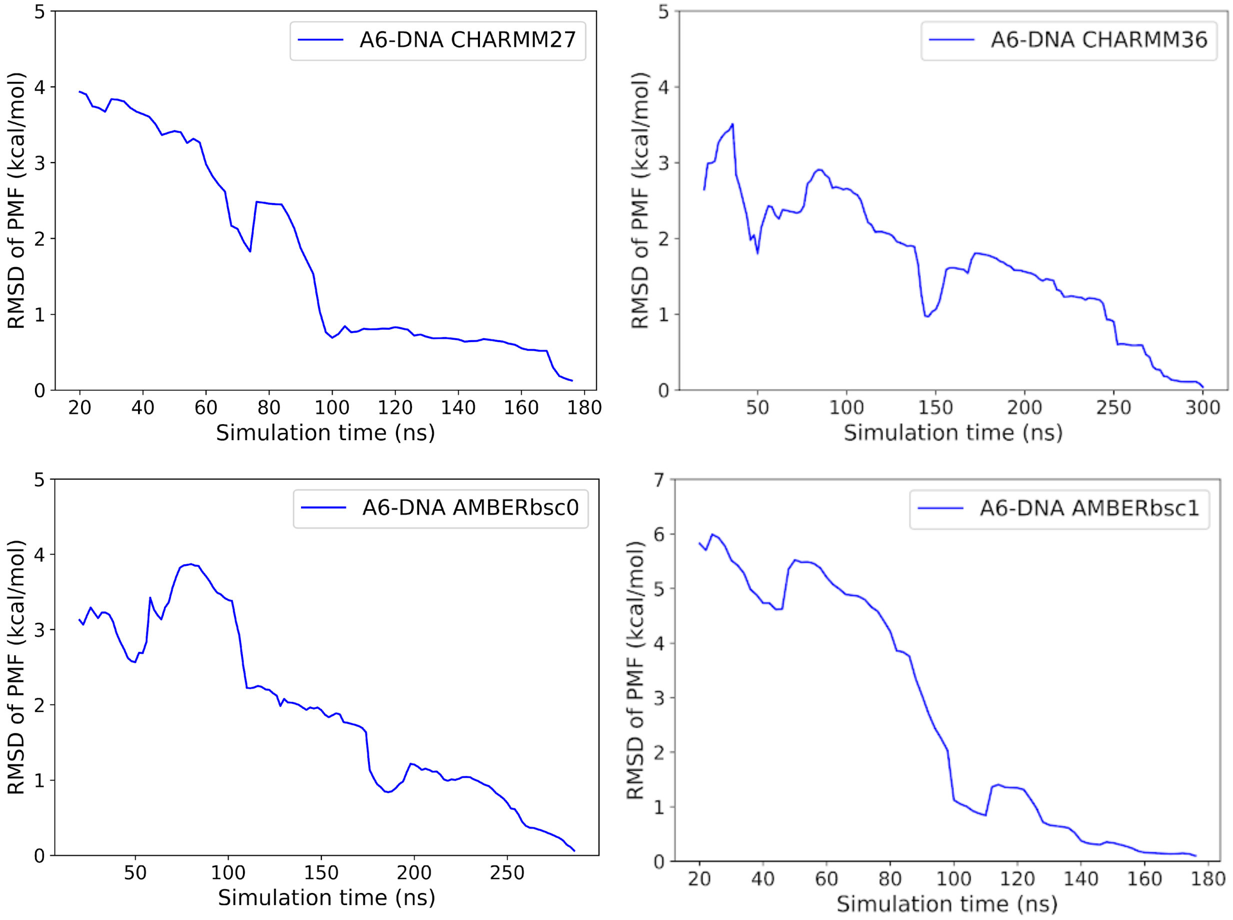


Figure S1: Root mean squared deviation of the 2D PMF with respect to the PMF at the end point of the meta-eABF simulation. PMFs were considered converged when the RMSD reaches 0.25 kcal/mol (following previous work[1]) and is less that 1 kcal/mol for the last 50 ns. For our meta-eABF simulations, CHARMM27 and AMBERbsc1 models converged within 200 ns, while CHARMM36 and AMBERbsc0 converged within 300 ns.

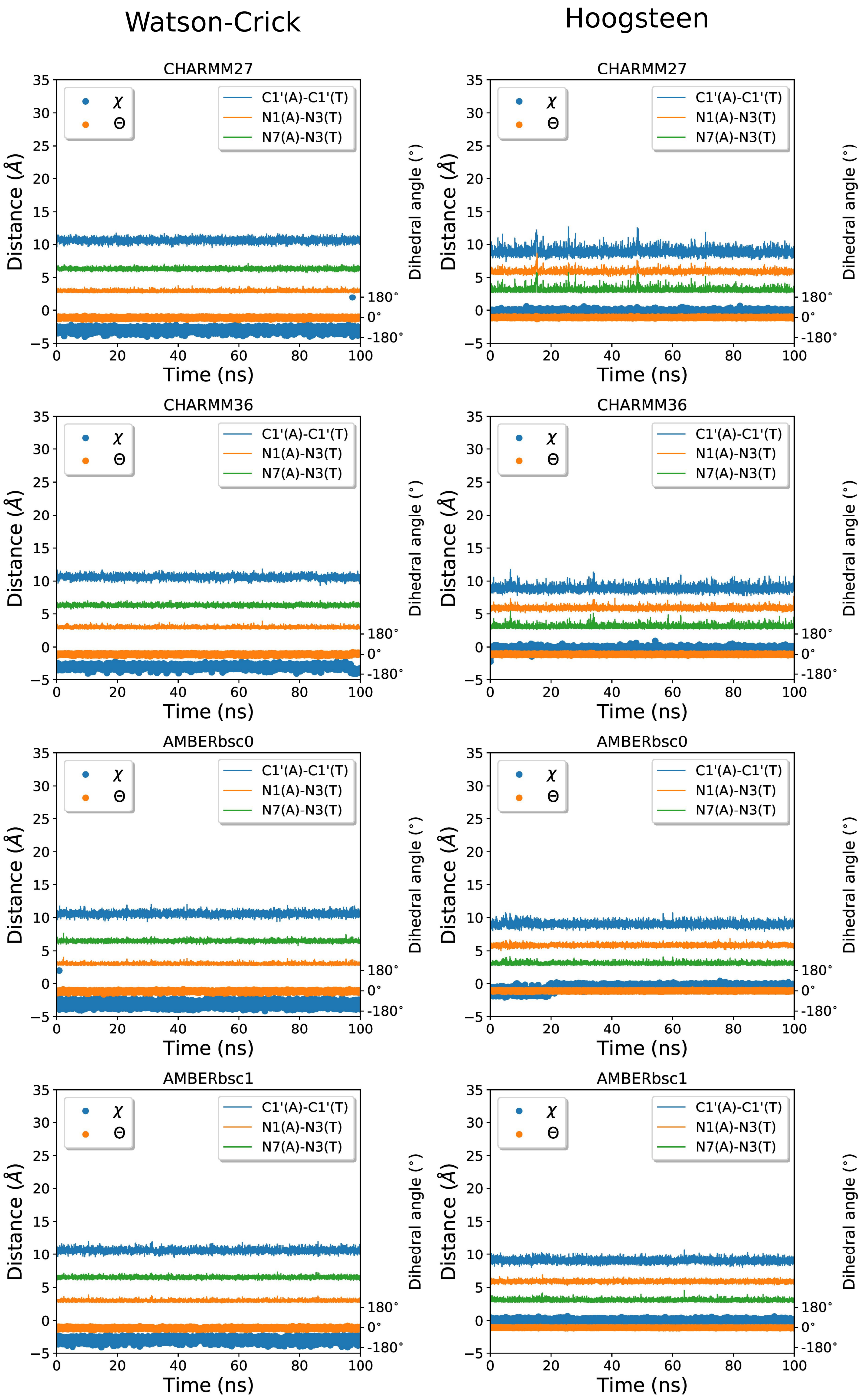


Figure S2: The hydrogen bond donor acceptor distances, the helix diameter measured as the C1'-C1' distance, and the two torsion angles $\chi$ and $\theta$ for the 100ns equilibrium simulations. As no transitions between WC and HG and no base pair melting is observed, we conclude that these two base pairing forms correspond to stable free energy minima in all four force field models.

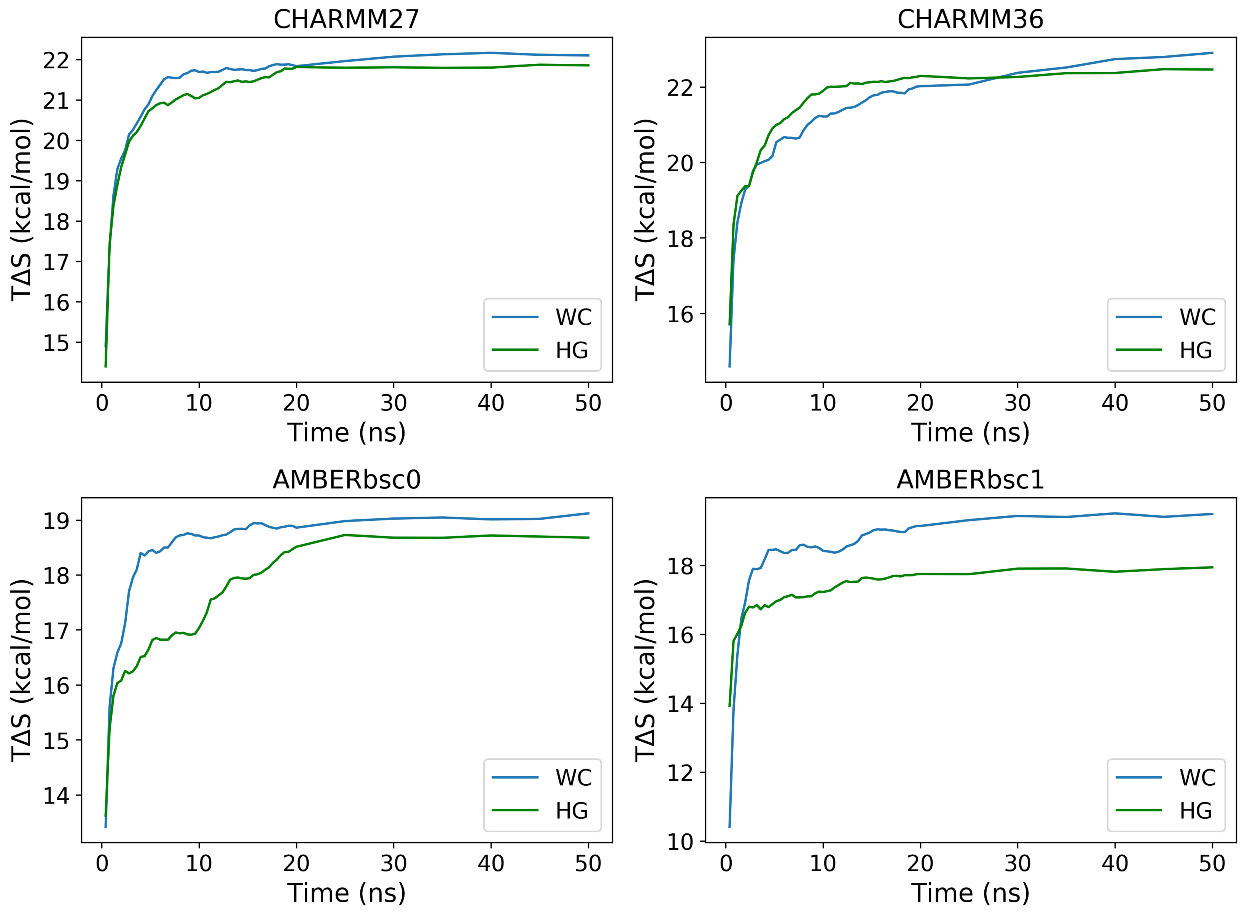


Figure S3: Convergence of quasi-harmonic entropy for different force fields. Only the 5th segment (50ns-100ns) of the 5 segments of the equilibrium trajectory has been used for calculating the convergence.

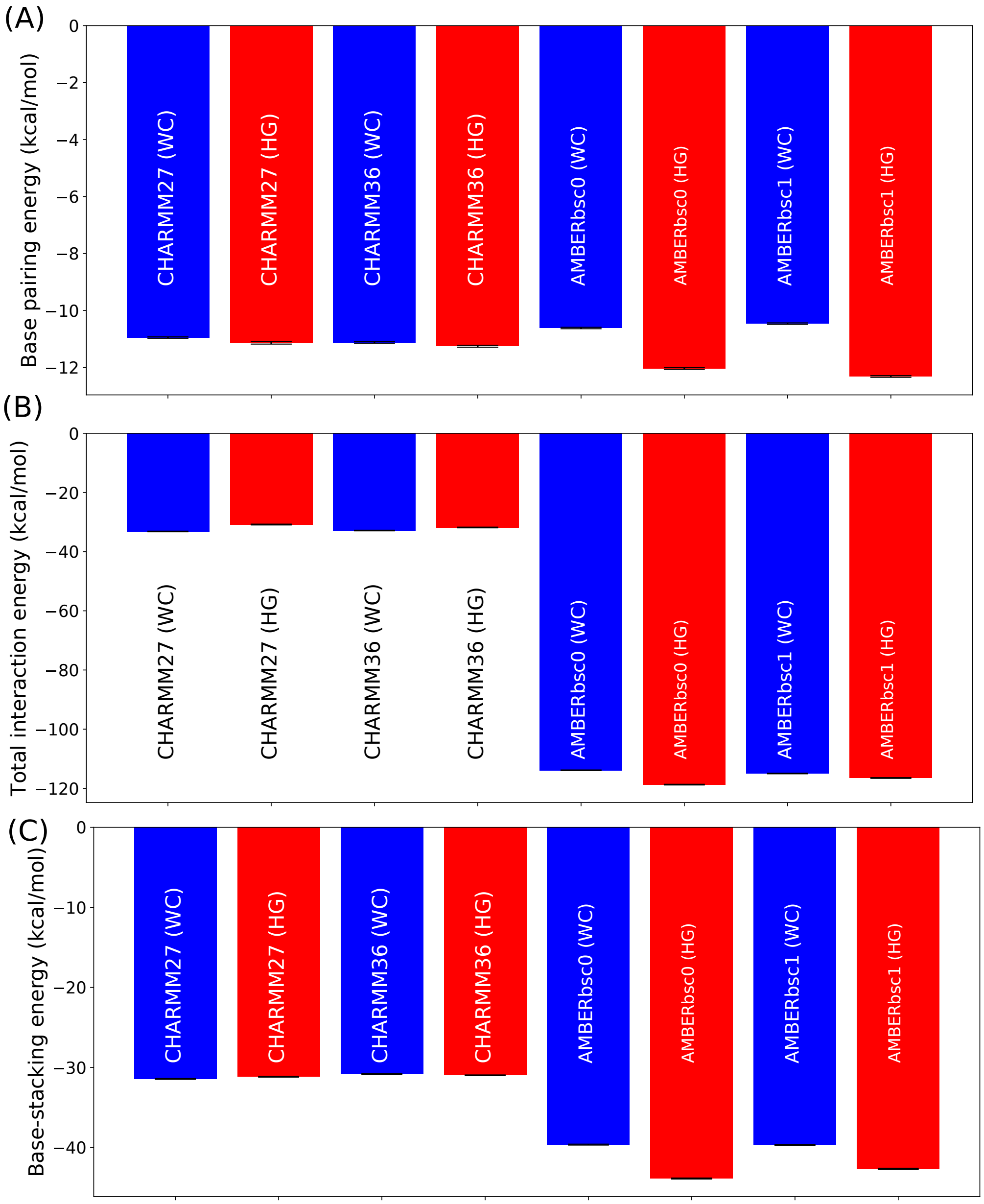


Figure S4: The absolute values of the following interaction energies in the A6-DNA: (A) between the A16 and T9 bases, (B) between the base pair with the rest of the DNA duplex, and (C) the stacking energy defined as the interaction energy between the $\pi$ conjugated region of the base pair with the two adjacent base pairs.

Table 1: The absolute and relative conformational entropy (in kcal/mol) of the A6-DNA duplex for the four different force fields.

| Force field | Watson Crick | Hoogsteen | HG - WC |
|---|---|---|---|
| CHARMM27 | $22.02 \pm 0.09$ | $21.82 \pm 0.09$ | $-0.20 \pm 0.13$ |
| CHARMM36 | $22.71 \pm 0.17$ | $22.28 \pm 0.14$ | $-0.44 \pm 0.22$ |
| AMBERbsc0 | $19.19 \pm 0.15$ | $18.46 \pm 0.25$ | $-0.73 \pm 0.29$ |
| AMBERbsc1 | $19.57 \pm 0.08$ | $18.19 \pm 0.53$ | $-1.37 \pm 0.54$ |

Table 2: The absolute and relative base pairing energies (in kcal/mol) of the A16-T9 base pair for the four different force fields.

| Force field | Watson Crick | Hoogsteen | HG - WC |
|---|---|---|---|
| CHARMM27 | $-10.95 \pm 0.03$ | $-11.14 \pm 0.04$ | $-0.19 \pm 0.05$ |
| CHARMM36 | $-11.12 \pm 0.03$ | $-11.26 \pm 0.03$ | $-0.13 \pm 0.04$ |
| AMBERbsc0 | $-10.62 \pm 0.03$ | $-12.04 \pm 0.03$ | $-1.42 \pm 0.04$ |
| AMBERbsc1 | $-10.46 \pm 0.02$ | $-12.32 \pm 0.03$ | $-1.86 \pm 0.04$ |

Table 3: The absolute and relative interaction energies (in kcal/mol) of the A16-T9 base pair with the rest of the DNA duplex for the four different force fields.

| Force field | Watson Crick | Hoogsteen | HG - WC |
|---|---|---|---|
| CHARMM27 | $-33.18 \pm 0.11$ | $-30.85 \pm 0.14$ | $2.34 \pm 0.17$ |
| CHARMM36 | $-32.87 \pm 0.11$ | $-31.84 \pm 0.12$ | $1.02 \pm 0.16$ |
| AMBERbsc0 | $-113.89 \pm 0.11$ | $-118.74 \pm 0.12$ | $-4.85 \pm 0.16$ |
| AMBERbsc1 | $-114.98 \pm 0.10$ | $-116.49 \pm 0.13$ | $-1.52 \pm 0.17$ |

Table 4: The absolute and relative stacking energies (in kcal/mol) of the A16-T9 base pair for the four different force fields.

| Force field | Watson Crick | Hoogsteen | HG - WC |
|---|---|---|---|
| CHARMM27 | $-31.43 \pm 0.05$ | $-31.16 \pm 0.05$ | $0.27 \pm 0.07$ |
| CHARMM36 | $-30.83 \pm 0.05$ | $-30.98 \pm 0.05$ | $-0.15 \pm 0.07$ |
| AMBERbsc0 | $-39.64 \pm 0.06$ | $-43.88 \pm 0.06$ | $-4.24 \pm 0.08$ |
| AMBERbsc1 | $-39.66 \pm 0.06$ | $-42.66 \pm 0.07$ | $-2.99 \pm 0.09$ |
